# Rof is a key regulator of Rho and transcription termination in bacteria

**DOI:** 10.1101/2023.09.01.555900

**Authors:** Jing Zhang, Shuo Zhang, Wei Zhou, Xiang Zhang, Guanjin Li, Luoxuan Li, Xingyu Lin, Fang Liu, Yue Gao, Zhenguo Chen, Yanjie Chao, Chengyuan Wang

**Author notes:** Equal contributions (Jing Zhang, Shuo Zhang, Wei Zhou, Xiang Zhang).

## Abstract

Transcription is crucial for the expression of genetic information and its efficient and accurate termination is required for all living organisms. Rho-dependent termination could rapidly terminate unwanted premature RNAs and play important roles in bacterial adaptation to changing environments. Although Rho has been discovered for about five decades, the regulation mechanisms of Rho-dependent termination are still not fully elucidated. Here we report the cryogenic electron microscopy structure of Rho-Rof antitermination complex. The structure shows that Rof binds to the open-ring Rho hexamer and inhibits the initiation of Rho-dependent termination. Rof’s N-terminal α-helix is key in facilitating Rof-Rho interactions. Rof binds to Rho’s primary binding site (PBS) and excludes Rho from binding with PBS ligand RNA at the initiation step. Further in vivo assays in *Salmonella Typhimurium* show that Rof is required for virulence gene expression and host cell invasion, unveiling a novel physiological function of Rof and transcription termination in bacterial pathogenesis.

## Main Text

The Rho factor is a ring-shaped ATP-dependent hexameric helicase that is conserved throughout the bacterial kingdom^1–12^. Rho plays an indispensable role in transcriptional termination and gene regulation in bacteria^1–4^. Coordinated with transcription-translation coupling and mediated by the transcription elongation factor NusG, Rho modulates factor-dependent transcription termination and regulates the gene expression^13–18^. Rho also separates transcription units, represses xenogenic genes^19^, silences antisense RNAs^20^, removes stalled RNAP from DNA thus maintaining the genome stability^21^, and prevents the formation of R-loops^22^. In the model bacterium *Escherichia coli*, about half of the transcription events are terminated by Rho factors^20,21^.

Five decades of genetic and biochemical experiments indicate that Rho-dependent transcription termination involves a series of transient steps^1–12^. The Rho-dependent pre-termination complexes were only recently determined at atomic resolution using cryoEM, unveiling the molecular details how Rho initiates transcription termination^16^. As the crucial first step, Rho recognizes a long C-rich RNA sequence (Rho utilization site; *rut* site) through its PBS and bind with the *rut* site^1–4^. Upon binding, Rho undergoes conformational changes from open-ring state to a catalytically competent, close-ring state. In the final step, Rho performs ATP-hydrolysis-dependent 5’ −> 3’ translocation on mRNA, applying mechanical force to the transcription elongation complex (TEC) and triggering termination. Whereas the latter steps are facilitated by a number of structural and regulatory proteins such as NusG and NusA, whether and how the initial RNA binding step is regulated remains little understood.

Rof (Rho-off, also called YaeO) is the only *E. coli* host factor that directly interacts with Rho and inhibits the Rho-dependent transcription termination^23–25^. Rof was proposed to bind to Rho and inhibit the PBS ligand binding^24^. However, whether Rof forms a protein complex with Rho and the basic nature of the interactions have been elusive. The regulatory and physiological functions of Rof as a conserved host factor in bacteria are completely unknown.

Herein, we determined the atomic cryo-EM structure of Rho-Rof antitermination complex and revealed the molecular interactions between Rho and Rof. Together with our recently determined Rho-dependent pre-termination complexes, the new structure shows that Rof directly binds with Rho N-terminal domain (PBS site) and disrupts the interactions between PBS ligand RNA and Rho. Rof regulates the initiation of Rho-dependent termination and inhibits termination efficiency. Our *in vivo* assays further showed that Rof plays crucial roles in virulence regulation in the model bacterial pathogen *Salmonella enterica serovar* Typhimurium (henceforth *S.* Typhimurium). Deletion of *rof* significantly reduced the expression of multiple virulence factors that are required for *Salmonella* invasion of host cells.

### Structure of Rho-Rof antitermination complex

The Rof and Rho proteins were individually expressed in *E. coli* and purified separately. Gel filtration assays showed that Rof is monomeric while Rho is hexameric in solution (Supplementary Fig. 1a, b). We assembled the Rho-Rof antitermination complex by mixing the purified Rho and Rof by the ratio as 1:1. The assembled complex were further validated in gel filtration assay, and the results confirmed the extensive interactions between Rof and Rho (Supplementary Fig. 1c). We subjected the assembled complex to single-particle reconstruction cryo-EM study and determined a 2.8 Å resolution structure of the Rho-Rof complex (Fig. 1a, Supplementary Fig. 2 and Supplementary Table 1).

**Fig. 1.**
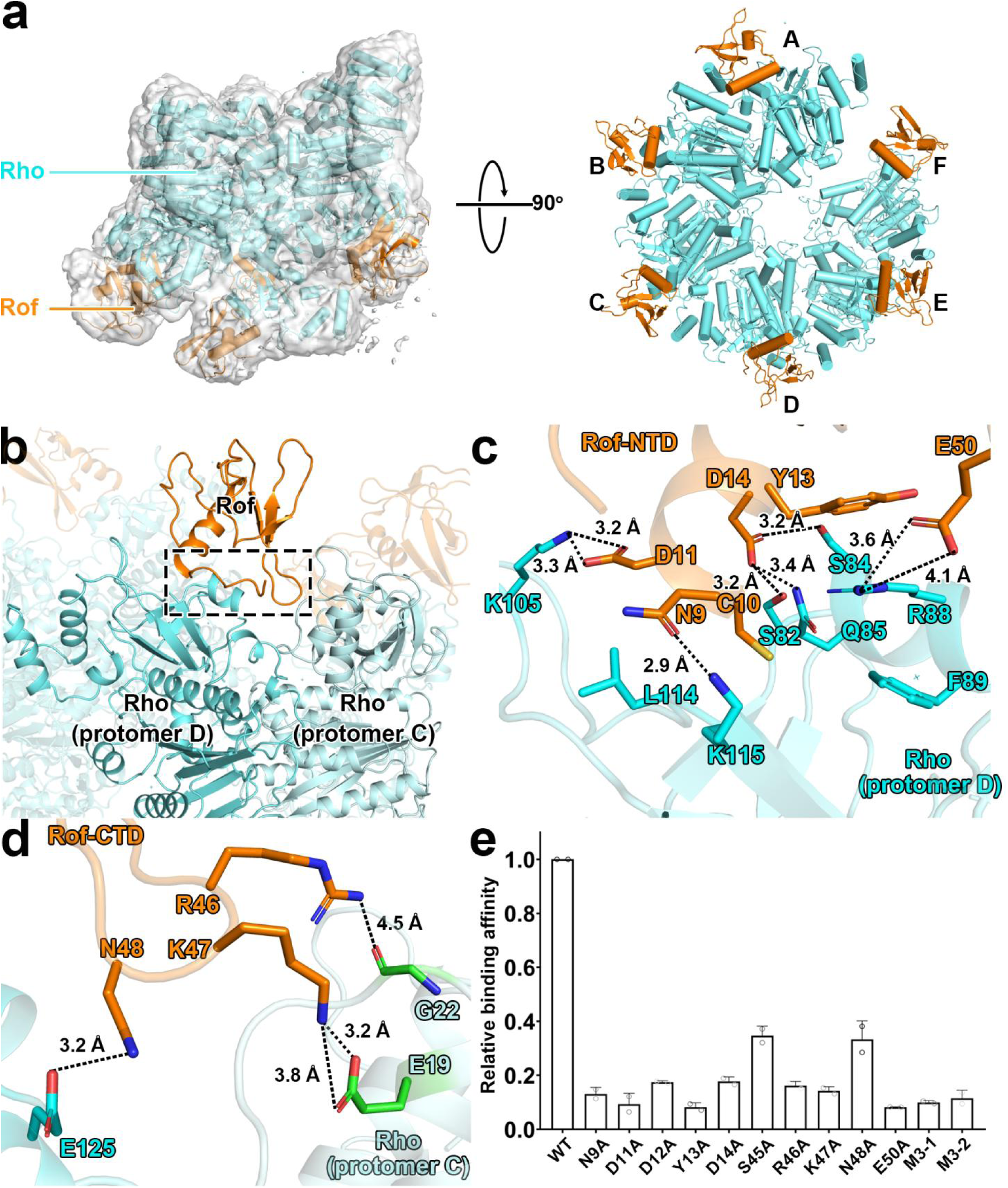
Structure of the Rho-Rof antitermination complex. (A) Cryo-EM structure of the Rof-Rho complex. Left panels, view orientation having the Rho hexamer central channel aligned with the y axis, showing the open-ring gap between Rho protomer A and protomer B. Right panels, orthogonal view orientation showing interactions of Rof with Rho protomers A-F. Images show EM density (gray surface) and fit (ribbons) for Rho. Rho and Rof are in cyan and orange. (B) Rho-Rof interactions in Rho-Rof antitermination complex. Overall interactions, focusing on Rof binding with Rho protomer D and C. The interface is highlighted by dashed black box. Rof, Rho (protomer D) and Rho (protomer C) are in orange, cyan, and light cyan. Other parts of the complex are shown in transparent color. (C) Details of interactions for Rof-NTD and Rho protomer D. Rof residues that interact with Rho are labeled and shown as orange sticks, Rho protomer D residues involving the interactions are shown as cyan sticks. Black dashed lines indicate H-bonds and salt bridges. (D) Details of interactions for Rof-CTD and Rho protomer D and C. Rho protomer C residues involving the interactions are shown as green sticks. Other colors and labels are as Fig. 1c. (E) Substitution of Rof residues involved in interactions with Rho decreased their in *vitro* binding affinity. Data for Biacore experiments are means of three technical replicates. Error bars represent ± SEM of n = 3 experiments. M3-1 represents N9A/D11A/D12A triple mutations, M3-2 represents R46A/K47A/N48A triple mutations.

The overall structure shows a Rho hexamer (open-ring state, comprising six protomers A-F) interacting with six Rof proteins in each of its protomers (Fig. 1a). The architecture of Rho is identical to previous reported open-ring state Rho structures, in which the midpoints of protomers A and F at either end of the ring are offset by 45 Å and leaves a 12 Å wide gap^11^. High-resolution data clearly show additional Rof density near the N-terminal of Rho with local resolution at ~4-5 Å (Fig. 1a and Supplementary Fig. 2). The densities of Rof in all protomers are clear except the one near protomer F, enabling unambiguous rigid-body docking of atomic structure of *Ec*Rof (PDB ID: 1SG5). The final models of Rof in the complex exhibit a conserved α-β-sandwich-like structure which was observed in both structures of *Ec*Rof and *Vc*Rof (PDB ID: 6JIE).

Rof shows conformational changes when interacting with Rho. The conformations of Rof structures in our model, particularly the orientation of the N-terminal α-helix (residues 10-21) differ from the NMR structure of *Ec*Rof^24^. However, they are more similar with the crystal structure of *Vc*Rof (RMSD 3.35 Å vs 3.78 Å, Supplementary Fig. 3a, b). These conformational changes are more likely caused by intermolecular interactions. As in the crystal structure of *Vc*Rof, two *Vc*Rof molecules interact with each other in the asymmetric unit via N-terminal α-helix^25^. In our cryo-EM structure, this α-helix is also involved in the interactions between Rof and Rho (Fig. 1b). The NMR structure of *Ec*Rof is monomeric and no protein-protein interactions have been observed^24^.

The N-terminal α-helix is crucial in facilitating interactions between Rof and Rho. In our cryo-EM structure, the spatial relationship between each Rof and the Rho protomers is identical. Rof binds extensively to Rho by attaching its N-terminal α-helix to the N-terminal of Rho (Fig. 1b and Supplementary Fig. 3c). The negative-charged residues Asn9, Asp11, Asp14 from Rof N-terminal α-helix form hydrogen bonds with Lys115, Lys105, Ser82, Gln85, Ser84 in Rho, respectively. In addition, residues Cys10 and Tyr13 stabilize the interface via van der Waals interactions with residues Leu114, Phe 89 (Fig. 1c).

The Rof’s β3-β4 loops (residues 45-50) make additional interactions with two Rho protomers. As showed in Fig. 1d, the residues Arg46, Lys47, Asn48, and Glu50 from β3-β4 loops make contacts with both protomer D and C by forming strong hydrogen bonds with residues Gly22, Glu19, Glu125 and Arg88, respectively (Fig. 1d).

All residues involved in the Rho-Rof interactions were verified through in vitro binding assays, confirming that Rof’s interactions with both protomers are crucial for its binding affinity (Fig. 1e). It appears that the open-ring gap between protomer F and A is causing a disruption in the interactions between Rof’s β3-β4 loops and Rho. As a result, the density of Rof in protomer F is considerably worse and has lower local resolution compared to the other protomers of Rho. The residues in the Rho-Rof binding site are conserved across different species, suggesting a general binding mechanism of protein interactions in bacteria (Supplementary Fig. 4).

### Antitermination mechanism of Rof in Rho-dependent termination

The crucial stage of Rho-dependent termination is the binding of the PBS ligand RNA with Rho^1–4^. Previously, we determined structures of Rho-dependent pre-termination complexes^16^. These complexes have NusG bridging Rho and RNAP, have mRNA tracing from RNAP transcription active center and threading through the central channel of Rho, and have 60 nucleotides of RNA interacting sequence-specifically with the exterior of Rho PBS (Fig. 2a, left panel). All six protomers of Rho hexamer interact with the PBS ligand RNA. Each Rho protomer is associated with ten nucleotides, in which 5 nt of the PBS ligand make potentially sequence-specific interactions with Rho protomer and another 5 nt connect the sequence-specifically recognized RNA segment between two protomers. In vitro transcription termination assay shows the PBS ligand is essential for subsequent steps in Rho-dependent termination. Without the PBS ligand, the termination efficiency of Rho-dependent termination is greatly decreased.

**Fig. 2.**
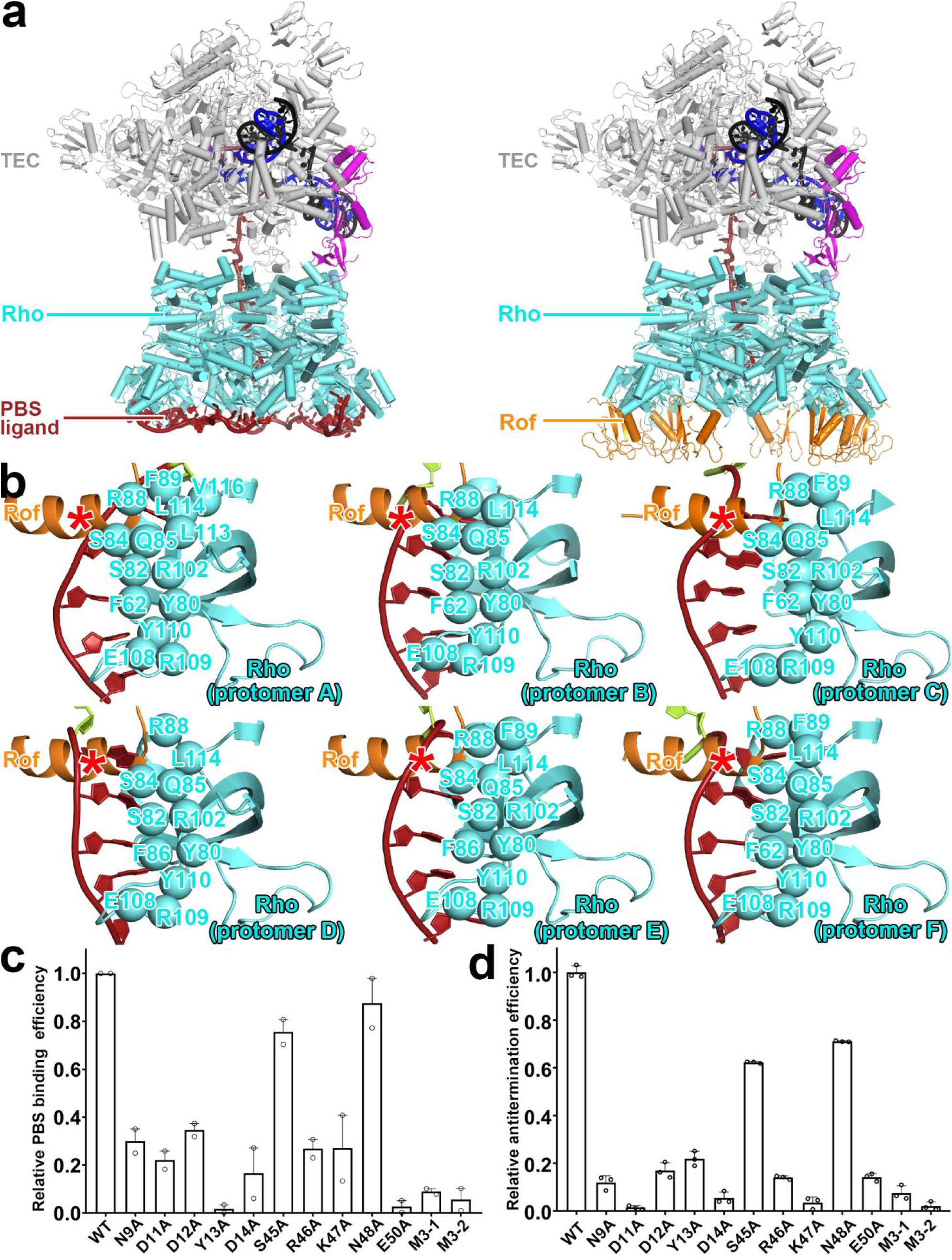
Anti-termination mechanism of Rof. **(A)** Superposition of Rho-Rof antitermination complex with Rho-dependent pre-termination complex. Left panel, overall structures of Rho-dependent pre-termination complex with λtR1 *rut* RNA (λtR1-Rho-NusG-TEC). Rho, cyan; RNAP in TEC, grey; non-template strand DNA, template strand DNA, and RNA, black, blue, and brick-red, respectively; NusG, magenta. Right panel, exactly same view with left panel, each Rof-Rho protomer are superposed with structure of λtR1-Rho-NusG-TEC, and replaced λtR1 *rut* RNA with Rof. Rof, orange; other colors and labels are as left panel. **(B)** Details of interactions for Rof/λtR1 *rut* RNA with Rho protomers A-F. Rho residues that interact with PBS-ligand RNA are shown as cyan spheres. λtR1 *rut* RNA is shown as brick-red; Rof N-terminal a helix is shown as orange; the competing binding site of Rof and PBS-ligand RNA is marked by red asterisk. **(C)** Relative inhibition efficiency of Rof on Rho-dC34 complex identified by electrophoretic mobility shift assay. Substitutions of residues involved in Rof-Rho interactions suppressed Rof’s binding affinity with Rho thus decreased Rof’s inhibition efficiency on Rho-dC34 formation. Data for in vitro transcription assays are means of three technical replicates. Error bars represent ± SEM of n = 3 experiments. M3-1 represents N9A/D11A/D12A triple mutations, M3-2 represents R46A/K47A/N48A triple mutations. **(D)** Relative *in vitro* transcription activity with Rho and Rof. Substitutions of residues involved in Rof-Rho interactions suppressed Rof’s antitermination efficiency on Rho-dependent termination thus suppressed in vitro transcription activity. Data for in vitro transcription assays are means of three technical replicates. Error bars represent ± SEM of n = 3 experiments. M3-1 represents N9A/D11A/D12A triple mutations, M3-2 represents R46A/K47A/N48A triple mutations.

Rof competes with the PBS ligand to bind to Rho and inhibits Rho-dependent termination. We superimposed each Rho protomer and corresponding Rof with the Rho-dependent pre-termination complex structure. The Rof binding sites in Rho are in the same location as the PBS ligand RNA according to the superposed model (Fig. 2a, right panel). Zoom-in view reveals the detail geometry of the conflict binding sites. In the Rho-dependent pre-termination complex, the PBS ligand RNA makes sequence-specific interactions with α-helix 4 and β-strands 4-5 of the Rho protomer. In this binding site, 5 nt of PBS-ligand RNA interact with a series of residues in each Rho protomer with a consensus sequence ((AACCC)_6_). The N-terminal α-helix of Rof makes extensive interactions with Rho and almost half of the residues (Ser82, Ser8e, Gln85, Arg88, Phe89, and Leu114) are interacting residues involved in the Rho-PBS ligand RNA binding. Molecular modeling suggests that, if α-helix of Rof were present, it would clash with PBS-ligand RNA (Fig. 2b).

Electrophoretic Mobility Shift Assay (EMSA) confirms that Rof could compete with PBS-ligand RNA, and disrupt the interactions between PBS ligand and Rho (Fig. 2c, Supplementary Fig 5). Mutations in Rof’s interacting interface could reduce or eliminate binding with Rho and promote PBS ligand binding. In vitro transcription assays confirm that the antitermination efficiency of Rof is depending on its inhibition efficiency for PBS ligand (Fig. 2c and d).

### Rof regulates the virulence gene expression in *Salmonella Typhimurium*

Given the indispensable role of Rho in bacteria, we asked whether Rof plays any physiological or regulatory roles *in vivo*. We first knocked out *rof* in the genomes of *E. coli* and *S. Typhimurium*, two well-established model organisms to study bacterial physiology and pathogenesis, respectively. Whereas the Δ*rof* mutants did not display any detectable phenotype in growth, overexpression of Rof inhibited *E. coli* growth under osmolarity stress (Figure 3a) and completely inhibited *Salmonella* growth and colony formation in standard LB medium (Figure 3b). The growth inhibition of *E. coli* was relieved by introducing single point mutations or triple mutations in Rof, suggesting that its regulatory function is dependent on the binding with Rho.

**Fig. 3.**
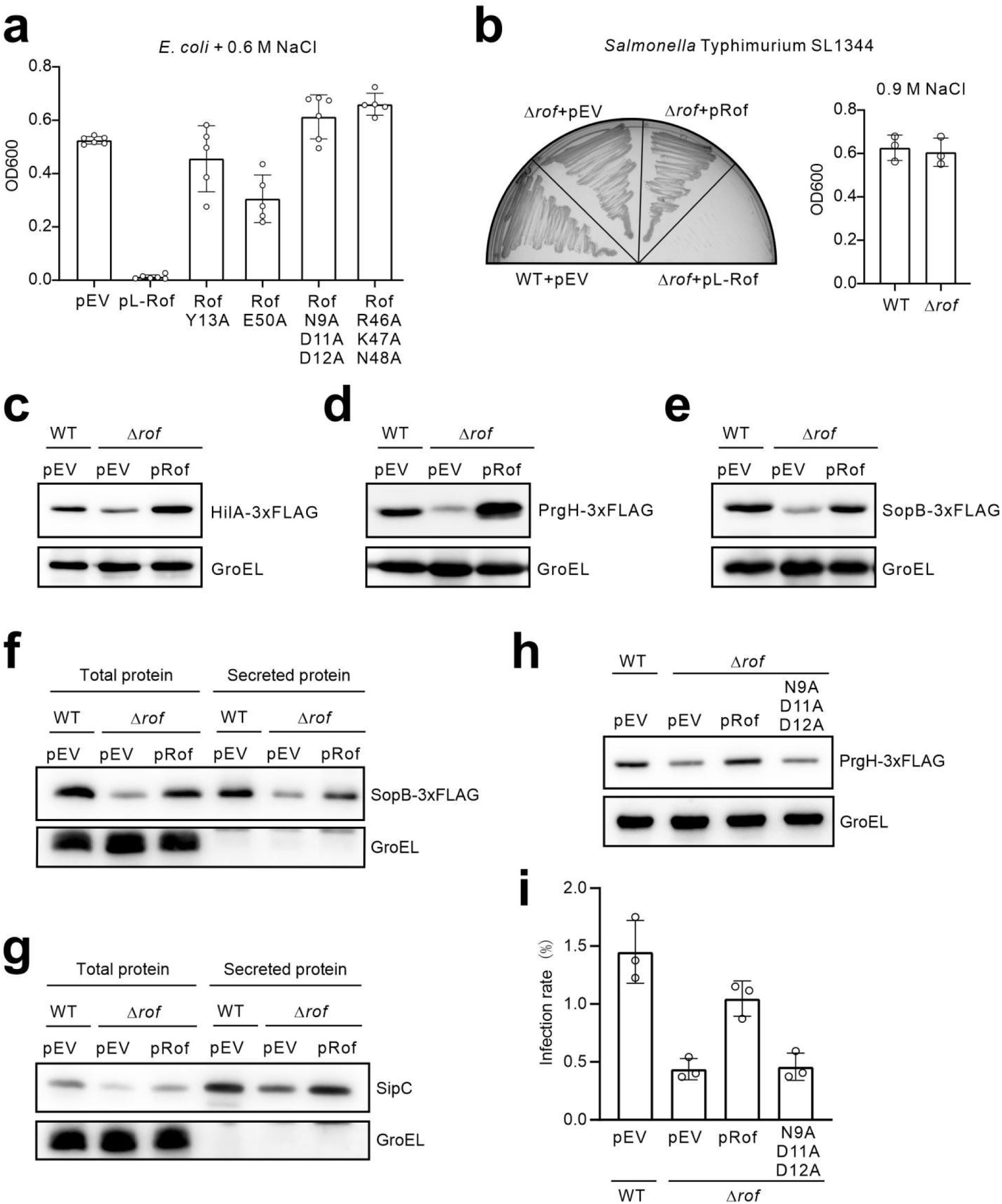
Rof regulates the virulence program of *Salmonella typhimurium*. **(A)** Rof inhibit the growth of *E. coli* under osmolarity stress. WT and mutant Rof were overexpressed from pET22b vectors. *E. coli* BL21 containing plasmids were grown overnight in LB supplemented with 0.6 M NaCl and 0.5 mM IPTG, and the final OD was recorded. pEV, empty vector. Error bars represent ± SD of n = 3 experiments. **(B)** Rof inhibit *Salmonella* growth and colony formation on LB. *Salmonella* WT and Δ*rof* mutant were transformed with pL-Rof plasmids, the resulting transformants were streaked on LB agar. No viable colony was observed for bacteria with pL-Rof, in which *rof* is driven by a constitutive P_LlacO_ promoter. pEV, empty vector. **(C)** Western blotting analysis of HilA-3xFLAG protein levels in WT *Salmonella* and Δ*rof* mutants. Total proteins were separated by 12% SDS-PAEG and analyzed using anti-FLAG antibody. GroEL served as a loading control. pEV, empty vector. pRof, Rof complementation plasmid. The *rof* gene with its own promoter was cloned into pZE12. **(D)** Western blotting analysis of PrgH-3xFLAG protein levels. **(E)** Western blotting analysis of SopB-3xFLAG protein levels. **(F)** Western blotting analysis of total proteins and secreted proteins in *Salmonella*. The secreted proteins were extracted from the culture supernatant. SopB was detected using anti-FLAG antibody; SipC was detected using anti-SipC antiserum. GroEL served as a non-secreted cellular protein control. (H) Western blotting analysis of PrgH-3xFLAG protein levels. (I) Rof is required for *Salmonella* invasion of epithelial cells. Human HCT-116 cell line was incubated with WT *Salmonella* and mutants using a MOI of 10. Gentamicin was added to kill extracellular bacteria after 40 min post-infection. The intracellular bacteria were harvested and enumerated after serial dilution and plating on LB agar. Error bars represent ± SD of n = 3 experiments.

Interestingly, a recent study has shown that Rho regulates the SPI-1 virulence program of *Salmonella*^26^, which is induced under the high osmolarity condition with low oxygen^27^ (similar as the gastrointestinal environment). We hypothesized that Rof may regulate virulence gene expression in *Salmonella*. To address this hypothesis, we introduced chromosomal 3× FLAGs to several representative genes in the SPI-1 regulatory cascade and examined their protein levels by Western blotting. Three genes were tagged, including *hilA* as the master transcriptional regulator of SPI-1 genes, *prgH* as a structural gene encoding part of the type 3 secretion system (T3SS), and *sopB* encoding an effector protein secreted via the T3SS^28^. When growing under the SPI-1 inducing condition (0.3 M NaCl + low oxygen), the Δ*rof* mutant indeed expressed lower amounts of HilA, PrgH and SopB proteins (Figure 3 c-e). The reduced levels in Δ*rof* were fully complemented by the *rof* gene provided *in trans*, confirming that Rof positively regulates the SPI-1 virulence gene expression in *Salmonella*. Not surprisingly, the secretion of SopB-3xFLAG and another untagged effector SipC, were reduced in the Δ*rof* mutant (Figure 3 f-g). The regulation of SPI-1 virulence program is dependent on the Rof-Rho complex formation, since the *rof* triple mutation disrupting the Rho-binding site failed to complement the Δ*rof* mutant (Figure 3h). The SPI-1 virulence program is required for Salmonella to invade host cells^27,28^. Using a human colon epithelial cell line HCT116, we have demonstrated that the Δ*rof* mutant has a reduced ability to invade host cells. This reduction was rescued by the WT *rof* gene on plasmid but not by the *rof* triple mutant, highlighting the regulatory function of Rof in virulence control in bacterial pathogenesis.

## Discussion

Our work shows the first structure of antitermination complex in Rho-dependent termination. The structure elucidates the basic nature of the interactions between Rof and Rho as a bipartite protein complex, in which the N-terminal α-helix of Rof is crucial in facilitating interactions and have conformational changes during the interactions. The structure clarifies the antitermination mechanism of Rof, showing that Rof competes with the PBS ligand to bind to Rho. Thus, Rof inhibits the initiation step of Rho-dependent termination. Our *in vivo* assays further illustrate the regulatory function of Rof in virulence control in bacterial pathogenesis. Based on these results and recent studies^26^, we suggest a model of Rof-Rho in virulence regulation in *S.* Typhimurium. Rho recognizes *rut* site sequence of target genes, initiates the termination of virulence genes and prevent their wasteful expression (Fig. 4, upper panel). During infection, Rof blocks Rho binding with *rut* site and inhibits Rho-dependent termination in virulence loci, leading to continued transcription of SPI-1 virulence genes and successful host invasion (Fig. 4, lower panel). To the best of our knowledge, these results for the first time showed a physiological role of Rof in virulence regulation in bacterial pathogens.

**Fig. 4.**
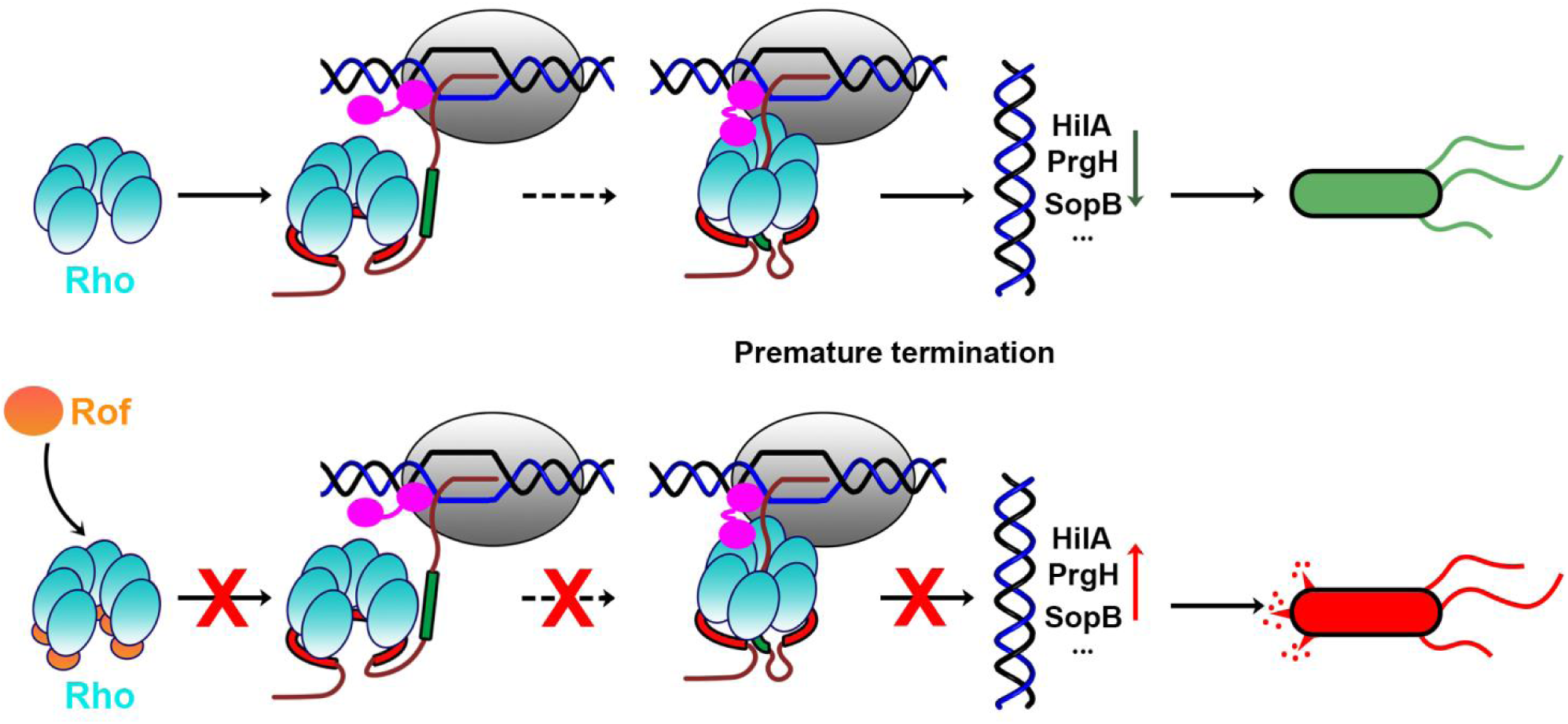
The proposed model of virulence gene regulation in *Salmonella* Typhimurium. Proposed regulatory mechanism of Rof and Rho in virulence gene expression in *Salmonella* Typhimurium. Upper panel (left to right). In the first step, the Rho hexamer in an open-ring state recognizes a long C-rich RNA sequence (PBS ligand; also known as *rut* site) through its primary binding site (PBS) on the exterior of the Rho hexamer. After a series of steps, Rho binds with short pyrimidine-rich RNA sequence (SBS ligand), changes from open-ring state to close-ring state, performs ATP-dependent 5’-3’ translocation on RNA towards the TEC, and terminates premature transcripts, and thus down-regulates the expression of *hilA, prgH, and sopB.* Rho, cyan; RNAP in TEC, grey; nontemplatestrand DNA, template-strand DNA, and RNA, black, blue, and brick-red, respectively; PBS ligand, red; SBS ligand, green; RNA 5′ to PBS ligand, RNA between PBS ligand and SBS ligand, and spacer RNA between SBS ligand and TEC, brick-red; NusG, magenta. Lower panel (left to right). Rof binds with Rho in its primary binding site, blocking PBS ligand RNA to bind Rho, thus prevents further transcription termination. The virulence genes *hilA, prgH, and sopB* are expressed without the inhibition of Rho-dependent termination.

Rof is a widely conserved protein in the phylum of Pseudomonadota (Proteobacteria). It is particularly abundant in the enterobacterial family, which contains many species of bacterial pathogens. (Supplementary Fig. 6). Phylogenetic analysis suggested that *rof* emerged later than *rho* in some bacterial lineages (Supplementary Fig. 7), suggesting that the *rof* gene may have evolved in response to complex environments, such as the intricate infection process in host cells. Besides Rof, other factors might also exist to regulate Rho-dependent termination in bacteria. Searching homologous structures of Rof using DALI service identified he Sm-like RNA-binding protein Hfq on the highest rank. Hfq could also be co-purified with Rho, and showed antitermination function in Rho-dependent termination^29,30^. This suggests that Hfq might be a structural homolog of Rof acting similarly on Rho, despite the low similarity in primary sequence (Supplementary Fig. 8). It remains to be clarified whether Rof and Hfq are the results of convergent evolution, and whether Hfq also inhibits the initiation step of Rho-dependent termination.

The initiation of Rho-dependent termination requires the transcription factor NusG, which bridges RNAP and Rho^1–4^. It has been shown that the interaction of NusG with Rho facilitates the formation of the catalytically competent, closed-ring state of Rho hexamer, triggering transcription termination^1^. Our work suggests that NusG could rescue the inhibitory effect of Rof at the initiation step. By performing *in vitro* transcription assays with NusG, we observed that the inhibition efficiency of Rof on Rho-dependent termination was reduced by the addition of NusG. As shown in Supplementary Fig. 9, Rof fully inhibit Rho’s termination while NusG could partly restore the termination activity. Our structures indicate no direct interaction between Rof and NusG, with Rof binding to Rho’s N-terminal domain while NusG binding to Rho’s C-terminal domain. It suggests that NusG does not relieve the inhibition of Rof on PBS ligand binding. Based on these results, we proposed that the initiation of Rho-dependent termination has two distinct mechanisms: one is PBS ligand dependent initiation which could be regulated by Rof, and the other is PBS ligand independent initiation which is mediated by NusG. It would be interesting to further dissect these two pathways of Rho-dependent termination and their respective regulatory targets and functions in bacteria in the future.

## Methods

### Protein purification

Rho was purified from BL21(DE3) cells carrying pET24b-rho as described previously. 2L cell culture was grown at 37°C in LB medium supplemented by kanamycin to OD_600_ = 0.8, induced with 1 mM IPTG at 18℃ for 20 hours, and harvested. Cells were resuspended in a lysis buffer (20 mM Tris-HCl, pH 7.6, 50 mM NaCl, 1 mM EDTA, 1 mM β-mercaptoethanol) and lysed by sonication on wet ice. Cell lysate was cleared by centrifugation for 30 min at 20,000g at 4°C, and applied to 5 ml HiTrap Q HP column (GE Healthcare) equilibrated with TGE buffer containing 50 mM NaCl and 1 mM β-mercaptoethanol. Rho was eluted by a 0.05-0.5 M NaCl gradient over 30 column volumes, concentrated, and diluted with TGE buffer containing 1 mM β-mercaptoethanol to adjust NaCl concentration to 0.05 M. Rho was next applied to 5 ml HiTrap HP heparin column (GE Healthcare) equilibrated with TGE buffer containing 50 mM NaCl and 1 mM β-mercaptoethanol. Rho was eluted by a 0.05-0.7 M NaCl gradient over 40 column volumes and concentrated to 3 ml volume. Finally, Rho was applied to 120 ml HiLoad 16/60 Superdex 200 size-exclusion column (GE Healthcare) equilibrated with 10 mM Tris-HCl, pH 7.6, 100 mM NaCl, 1 mM β-mercaptoethanol. Fractions containing pure Rho were combined and concentrated using 15 ml Amicon Ultra 100 kDa MWCO concentrators to 100 µM. The product (purity >98%) was stored in aliquots at −80°C.

Rof was purified from BL21(DE3) cells transformed with pET22b-rof. The protein expression was induced with 1 mM IPTG at 18 °C for 20 h at OD_600_ of 0.7. The cell pellet was lysed in lysis buffer (50 mM Tris-HCl, pH 7.6, 200 mM NaCl, 5% (v/v) glycerol, 0.1 mM PMSF) using an sonication. The supernatant was loaded on a 5 mL Ni-NTA column that was subsequently washed and eluted with lysis buffer B containing 200 mM imidazole. The eluted fractions were concentrated to 3 ml and applied to 120 ml HiLoad 16/60 Superdex 200 size-exclusion column (GE Healthcare) equilibrated with 10 mM Tris-HCl, pH 7.6, 100 mM NaCl, 1 mM β-mercaptoethanol. The fractions containing target proteins were concentrated to 4 mg/mL, and stored at −80 °C. Rof derivatives were prepared by the same procedure.

RNAP holo was prepared from BL21 Star (DE3) cells containing pIA900 (encodes *E. coli* RNAP β’ with C-terminal hexahistidine tag, β, α, and ω subunits) and pRSFDuet-rpoD plasmids. The product (purity >95%) was stored in aliquots in RNAP storage buffer (10 mM Tris-HCl, pH 7.6, 100 mM NaCl, 0.1 mM EDTA, and 5 mM dithiothreitol) at −80°C.

### Cryo-EM sample preparation

For cryo-EM grid preparation, samples (3 μL at a concentration of ~10.5 mg/mL) were applied to freshly glow-discharged Quantifoil R1.2/1.3 holey carbon grids. After incubation for 5 s at 6 °C and 100% humidity, the grids were blotted for 1 s with blot force 2 in a Thermo Fisher Scientific Vitrobot Mark IV and plunge-frozen in liquid ethane at liquid nitrogen temperature. The grids were prepared in the H2/O2 mixture for 60 s using a Gatan 950 Solarus plasma cleaning system with a power of 5 W. The ø 55/20 mm blotting paper is made by TED PELLA used for plunge freezing.

### Cryo-EM structure determination: data collection and data reduction

Cryo-EM data for Rho-Rof complex was collected at the Fudan University Cryo-EM core Facility, using a 300 kV Titan Krios (FEI/ThermoFisher) electron microscope equipped with a post-GIF Gatan K3 direct electron detector (Gatan). Data were collected automatically in the super-resolution mode, using Serial-EM with a nominal magnification of 105,000x, a calibrated pixel size of 0.595 Å/pixel on the image plane, and with defocus values ranging from −0.8 to −2.0 μm. Each micrograph stack was dose-fractionated to 40 frames with a total electron dose of ~50 e−/Å2 and a total exposure time of 3.6 s. 6033 micrographs of Rho-Rof complex were collected for further processing.

Data were processed as summarized in Figs. S1A-D. Data processing was performed using a Tensor TS4 Linux GPU workstation with four GTX 3090 graphic cards (NVIDIA). Dose weighting motion correction (3×3 tiles; b-factor = 150) were performed using Motioncor2^31^. Contrast-transfer-function (CTF) estimation was performed using CTFFIND-4.1^32^. Subsequent image processing was performed using Relion 3.0^33^. Automatic particle picking with Laplacian-of-Gaussian filtering yielded an initial set of 2,399,274 particles. Particles were extracted into 500×500 pixel boxes and subjected to rounds of reference-free 2D classification and removal of poorly populated classes, yielding a selected set of 1,099,028 particles. The selected set was auto-refined using a mask with a diameter of 300 Å, yielding a reconstruction at 3.3 Å overall resolution. The resulting 3D auto-refined particles were further done with post-processing with a soft masked, yielding reconstructions at 2.8 Å.

The initial atomic model for Rho-Rof complex was built by manual docking using open-ring Rho (PDB ID: 1PV4) and a crystal structure of *E. coli* Rof (PDB ID 1SG5) in UCSF Chimera^34^. For Rho (residues 280-284) and Rof (residues 1-5), density was absent, suggesting high segmental flexibility; these segments were not fitted. Refinement of the initial model was performed using real_space_refine under Phenix. The Rof and Rho were rigid-body refined against the map, followed by real-space refinement with geometry, rotamer, Ramachandran-plot, C β, non-crystallographic-symmetry, secondary-structure, and reference-structure (initial model as reference) restraints, followed by global minimization and local-rotamer fitting. Secondary-structure annotation was inspected and edited using UCSF Chimera. Rof were subjected to iterative cycles of model building and refinement in Coot^35^. The final atomic model at was deposited in the Electron Microscopy Data Bank (EMDB) and the Protein Data Bank (PDB) with accession codes EMDB 37342 and PDB 8W8D (Table S1).

### Electromobility Gel Shift Assays

Assays of nucleic acid binding to full-length Rho were carried out as described in previous report^36^. Chemically synthesized oligonucleotide dC34 (3 uM, final concentration), Rho (3 uM, final concentration) was mixed with Rof or its derivatives (6 uM, final concentration) in the reaction buffer (25 mM Tris-HCl, pH 8.0, 5 mM MgCl_2_, and 50 mM KCl, 10% glycerol). Bovine serum albumin (BSA, 200ug/ml) was used as a non-specific protein competitor. Each sample was incubated at 37℃ for 15 min and then analysed on a 6% polyacrylamide gel (acrylamide/bisacrylamide ratio of 29:1) in TBE (90 mM Tris, 44 mM boric acid, 2 mM EDTA) followed by SYBR-Gold staining. DNA bands were visualised using a gel documentation system (Tanon 2500R), and their intensities were quantified using ImageJ software (NIH Image, National Institutes of Health, Bethesda, MD, USA; online at: http://rsbweb.nih.gov/ij/).

### Fluorescence-detected In vitro transcription assay

The experiment was performed as in previous report^37^. Briefly, *E.coli* RNAP holo (final concentration: 50nM) was incubated with DNA template (final concentration: 16nM) in the reaction buffer at 37℃for 15min. Then, Rho (final concentration: 240nM) with or without YaeO (final concentration: 2.8 uM) was added and incubated at 37℃ for 10min. Finally, TOl-3PEG-Biotin (final concentration: 0.5 uM) was added and the reactions were initiated by addition of NTP mix (final concentration: 0.5 uM). The fluorescence signals were collected using SparkControl software on a plate reader (SPARK, TECAN) at an excitation wavelength of 510/9.5 nm and an emission wavelength of 550/9.5 nm.

### Biacore

BIAcoreT200 was used to study the direct binding of Rof and mutants to Rho. Rho was immobilized (Target: 1000 resonance units (RU), Real: 1176 RU) by amine group coupling on research grade CM5 sensor chips at pH 4.5. Rof and mutants in HEPES buffer (10 mM HEPES (pH 7.4), 300 mM NaCl, and 0.05% (v/v) Tween-20) were then injected over the immobilized Rho at 25℃. Kinetics constants (KD) were derived by Scatchard analysis or nonlinear curve fitting of the standard Langmuir binding isotherm using BIAevaluation version 4.1 (BIAcoreT200).

### *Salmonella* strains and growth conditions

*Salmonella enterica* serovar Typhimurium strain SL1344 was used as wild-type. Strains with deletions or chromosomally 3xFLAG epitope-tagging were constructed using the λ-Red recombinase method. For plasmid construction, the desired gene fragments were generated by PCR amplification using *S.* Typhimurium SL1344 or *E. coli* K-12 MG1655 genomic DNA as template and, after digestion with restriction enzymes, were cloned into the corresponding sites of the indicated vectors. Mutations and plasmid inserts were confirmed by sequencing.

*Salmonella* was routinely cultured at 37 ℃ on LB agar or with shaking at 220 rpm in liquid LB media (10 g/L tryptone, 5 g/L yeast extract, 5 g/L NaCl). Overnight cultures were grown from a single colony, diluted 1:100 in fresh medium, and grown to the indicated OD600. For SPI-1 condition culture, a single colony was inoculated into 15 ml LB containing 0.3 M NaCl in tightly sealed 15 ml Falcon tubes, incubated overnight at 37 ℃ without agitation. The following compounds were added at the following final concentrations when appropriate: IPTG at 0.5 mM; ampicillin (Amp) at 100 μg/mL; kanamycin (Kan) at 50 µg/ml, chloramphenicol (Cm) at 20 µg/ml; hygromycin (Hyg) at 100 µg/ml.

### Extraction of secretory proteins from culture medium

*Salmonella* was cultured overnight under the SPI-1 condition. 2 ml culture was centrifuged at 12000 rpm for 5 min at room temperature. 1.5 ml supernatant was carefully transferred into a new tube, and added with 0.25 vol of 50 % TCA (Trichloroacetic acid). The mixture was incubated at 4 ℃ for 1 h, then the protein pellet was collected by centrifugation at 12000 rpm at 4 ℃ for 30 min. The pellet was washed twice with ice-cold acetone, and air-dried completed until the acetone was evaporated. The protein pellet was finally dissolved in 20 μl 1× protein loading buffer.

### Western blotting analysis

Bacterial samples were resuspended in 1× protein loading buffer and boiled at 98 °C for 8 min. 0.05 OD of proteins samples were loaded each lane. Proteins were transferred onto PVDF membranes (#10600023, Cytiva) for 45 min at 150 V in transfer buffer. Membranes were blocked for 1 h at room temperature in 1× TBST buffer with 5% (w/v) skim milk (#A600669-0250, Sangon), washed with TBST twice and incubated with monoclonal α-FLAG (Sigma-Aldrich #F1804-5MG; 1:10,000), α-SipC (1: 3,000) or α-GroEL (Sigma-Aldrich #G6532; 1:10,000) antibodies for 1 h at room temperature. After three TBST washes, membranes were incubated with secondary α-mouse or α-rabbit HRP-linked antibodies (Sigma-Aldrich #A0168; 1:10,000 or Sangon #D110087-0100; 1:10,000) for 1 h at room temperature. Chemiluminescence was developed using the high sensitive ECL luminescence reagent (#C500044-0100, Sangon), and then visualized on ChemiDocTM XRS + and quantified using ImageLabTM Software.

### *Salmonella* infection assay

Human colon cancer cell line HCT116 was purchased from CCTCC (#SCSP-644, Shanghai, China). The cells were grown in high-glucose Dulbecco’s Modified Eagle’s (DMEM) Medium (#12491015, Gibco) supplemented with 10% fetal bovine serum (FBS) (#2534387, Gibco), 50 U/ml penicillin and 50 μg/ml streptomycin at 37 ℃ with 5% CO2 and humidified atmosphere. Before infection, 1×10^5^ HCT116 cells were seeded in 1 ml complete DMEM (twelve-well format). *Salmonella* was grown under the SPI-1 condition. Bacterial cells were harvested by centrifugation (2 min at 12,000 rpm, room temperature), washed twice with phosphate-buffered saline (PBS, #B548117-0500) and resuspended in DMEM. Infection of HCT116 cells was carried out by adding the bacterial suspension directly to each well, at a multiplicity of infection (MOI) of 10. Then cells were incubation for 40 min at 37 °C in 5% CO2 and humidified atmosphere. After washing twice with PBS, cells were added with medium containing 100 μg/ml gentamicin, and incubated for 1 h to kill extracellular bacteria. Afterwards, the infected cells were washed two times with PBS, and added with fresh DMEM containing 50 μg/ml gentamicin for 1 h. The infected cells were washed with PBS and lysed by the addition of PBS with 0.1 % Triton X-100 (#A600198-0500, Sangon). After serial dilution of the lysate in PBS, intracellular bacteria were enumerated on LB agar plates and incubated overnight at 37 ℃.

## Supplementary Figure legends

**Fig. S1.**
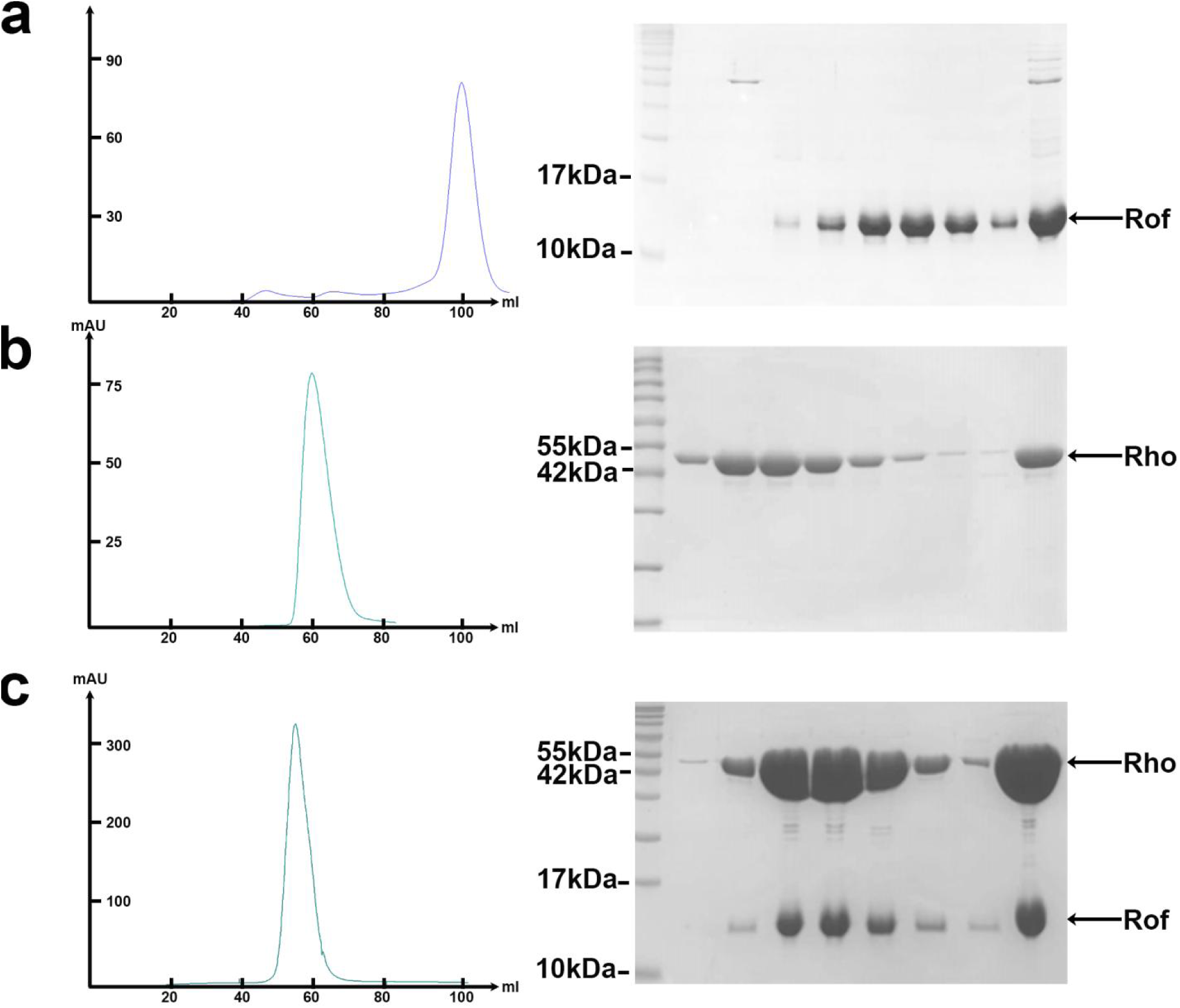
Purification and verification of *E. coli* Rof and Rho. (A) Left panel, Chromatogram map of gel filtration of *E. coli* Rof; right panel, SDS-PAGE of the purified *E. coli* Rof. (B) Left panel, Chromatogram map of gel filtration of *E. coli* Rho; right panel, SDS-PAGE of the purified *E. coli* Rho. (C) Left panel, Chromatogram map of gel filtration of *E. coli* Rho-Rof complex; right panel, SDS-PAGE of the purified *E. coli* Rho-Rof complex.

**Fig. S2.**
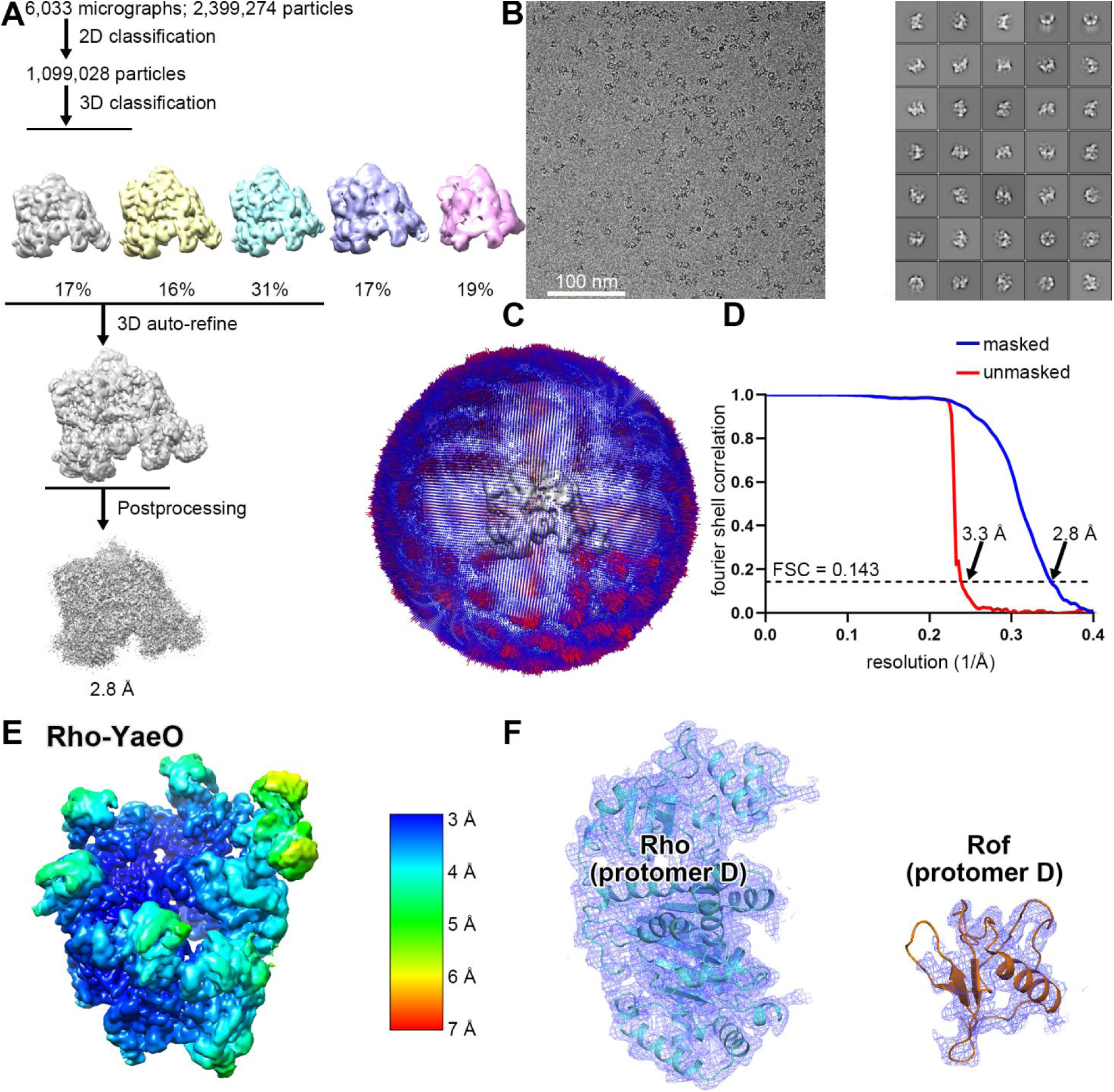
Structure determination of Rho-Rof antitermination complex. **(A)** Data processing scheme (Table S1). **(B)** Representative electron micrograph and 2D class averages (50 nm scale bar in right subpanel). **(C)** Fourier-shell-correlation (FSC) plot. **(D)** Orientation distribution. **(E)** EM density map coloured by local resolution (horizontal inversion view as in Fig. 2a, left). **(F)** Representative EM density (blue mesh) and fits (ribbons) for Rho and Rof.

**Fig. S3.**
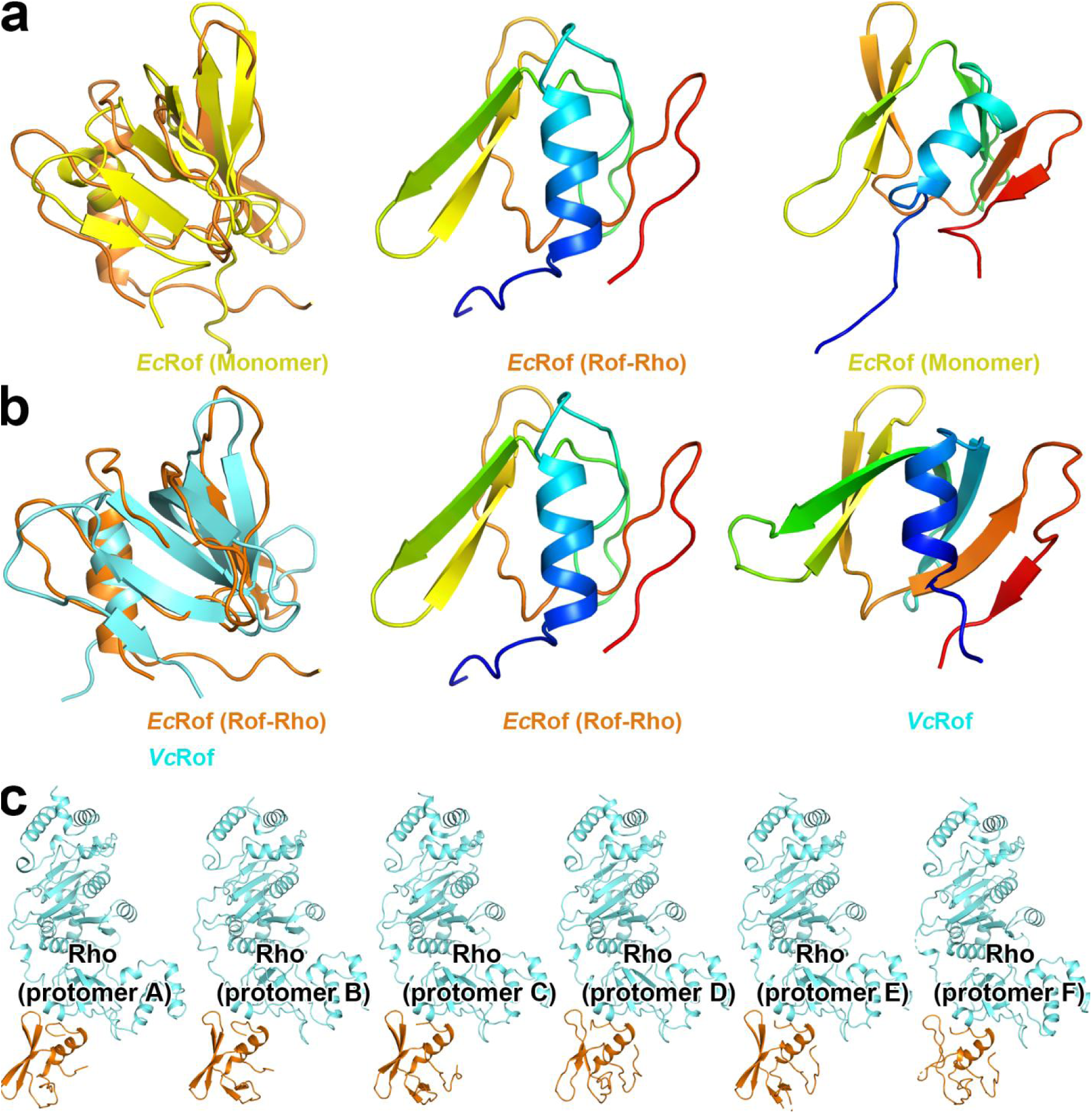
Structural comparison of Rof. **(A)** Superposition of *Ec*Rof. First panel, *Ec*Rof (this work) and *Ec*Rof (Monomer, determined by NMR method, PDB ID 1SG5) are shown in orange and yellow, respectively. Second panel, *Ec*Rof (this work) show in rainbow with N-terminal colored blue to C-terminal colored red; Third panel, *Ec*Rof (Monomer) colored as second panel. *Ec* is short for *Escherichia coli*. **(B)** Superposition of *Ec*Rof and *Vc*Rof (PDB ID 6JIE). First panel, *Ec*Rof (this work) and *Vc*Rof are shown in orange and cyan, respectively. *Vc* is short for *Vibrio cholerae*. **(C)** Structures of Rof-Rho in each protomers A-F.

**Fig. S4.**
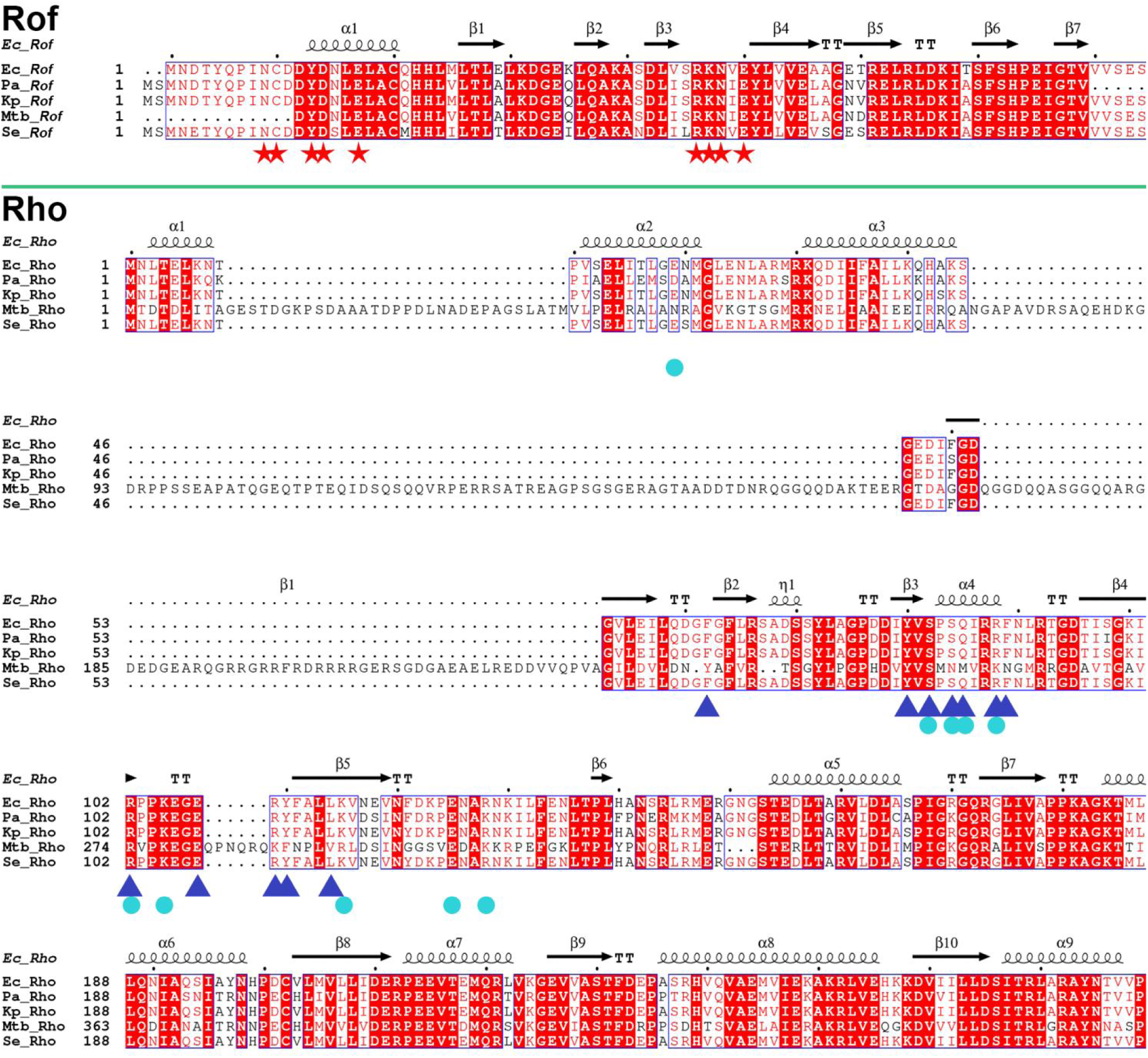
Sequence alignment of Rof and Rho. Upper panel, amino acid sequence alignment of Rof from *Ec, Escherichia_coli; Pa, Pseudomonas aeruginosa; Kp, Klebsiella pneumoniae; Se, Salmonella enterica; Mtb, Mycobacterium tuberculosis*. The residues of Rof-Rho interface from Rof are indicated by red stars. Lower panel, amino acid sequence alignment of Rof from *Ec, Escherichia_coli; Pa, Pseudomonas aeruginosa; Kp, Klebsiella pneumoniae; Se, Salmonella enterica; Mtb, Mycobacterium tuberculosis*. The residues of Rof-Rho interface from Rho are marked by cyan circles and residues of Rho-RNA interface from Rho are indicated by blue triangles.

**Fig. S5.**
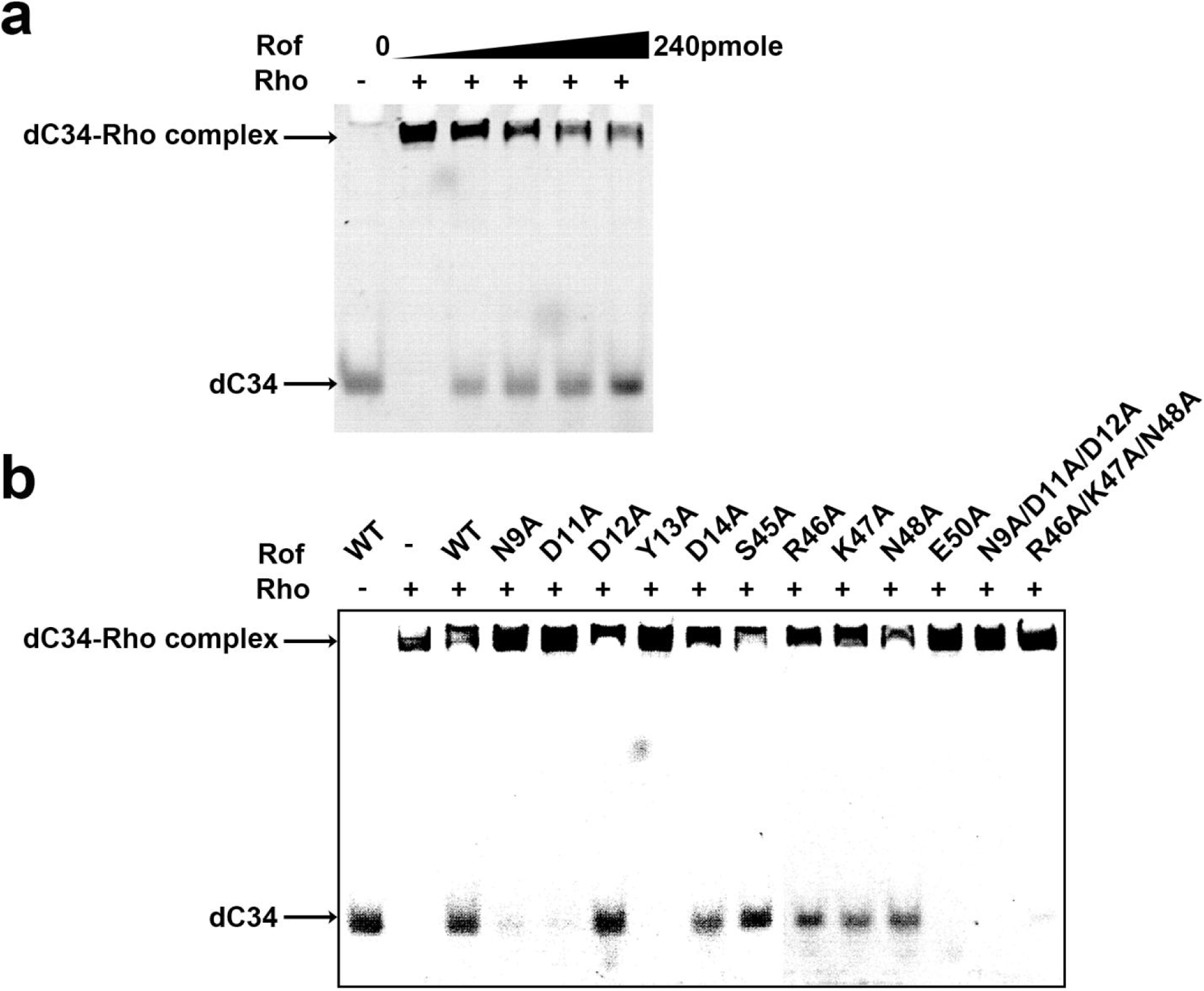
Electrophoretic mobility shift assay for Rho and PBS ligand RNA. **(A)** EMSA assay for Rho and dC34 binding with or without Rof. Rho and 3μm dC34 were added into the reaction system. Bands of dC34, dC34-Rho complex are indicated by arrows on the left, respectively. **(B)** EMSA assay for Rho and dC34 binding with or without Rof and its mutations. 3 uM Rho and 3 dC34 were added into the reaction system. Bands of dC34, dC34-Rho complex are indicated by arrows on the left, respectively.

**Fig. S6.**
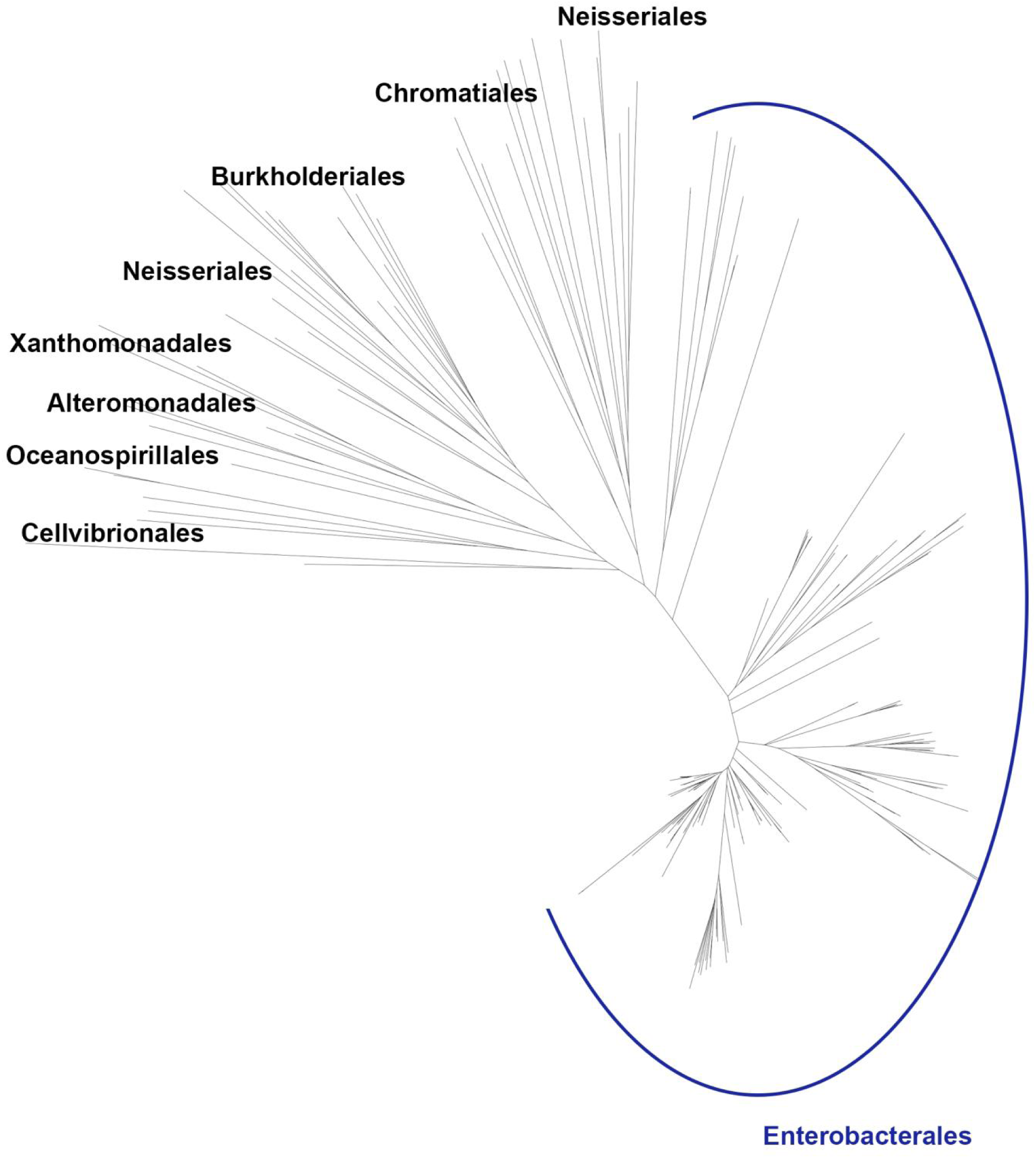
Phylogenetic analysis of the distribution of Rof in bacteria. 206 protein sequences of Rof are used in constructing the neighbor-joining phylogenetic tree. The family names of bacteria are shown on the top branch of the tree. The sequences belong to the Enterobacterales are highlighted with blue label and solid line.

**Fig. S7.**
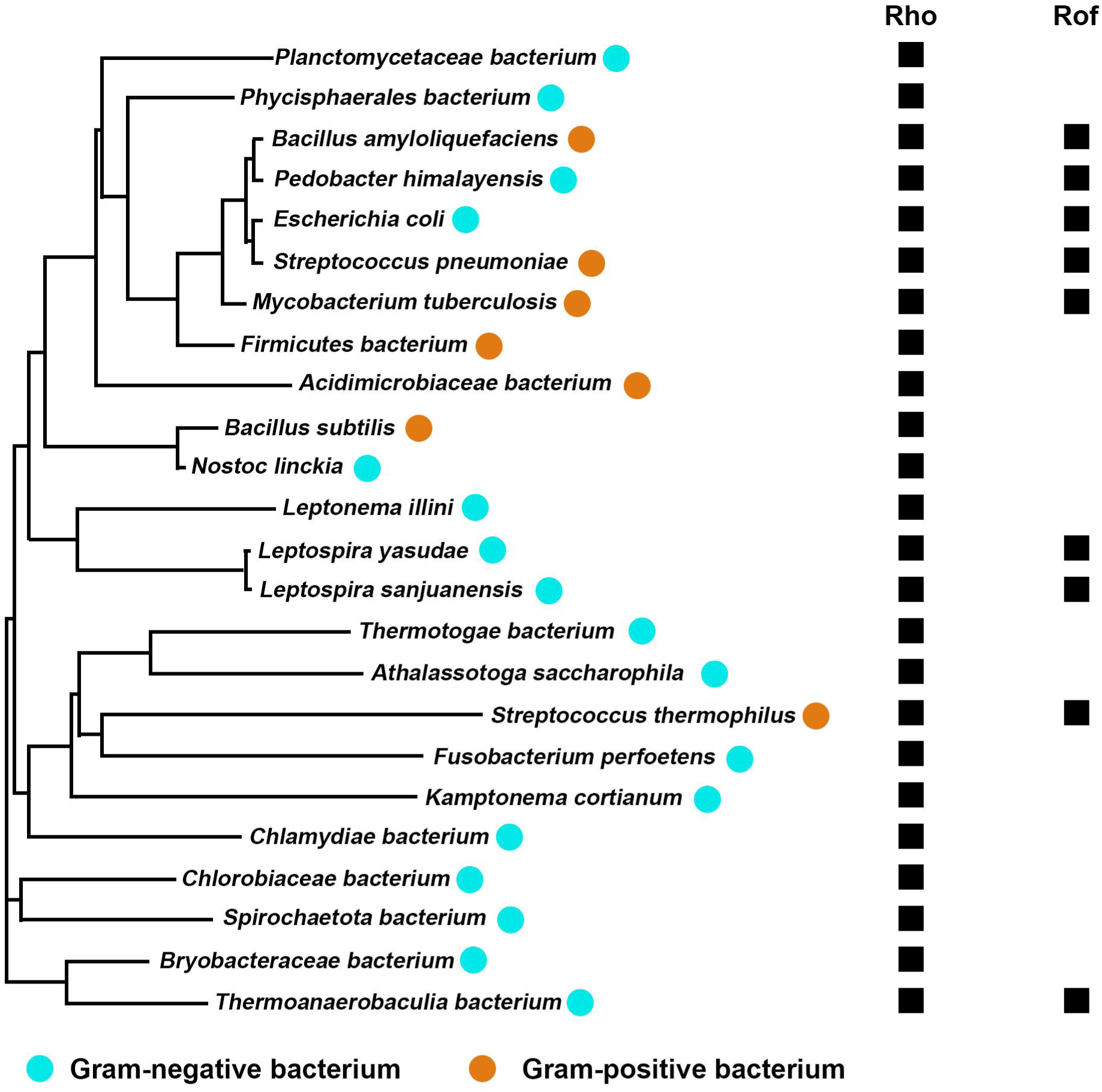
Phylogenetic analysis of the distribution of Rho and Rof in bacteria. Rho sequences from different species from Terrabacteria to α-proteobacteria are used in constructing the neighbor-joining phylogenetic tree. Gram-negative bacterium species are marked as cyan circles, and Gram-positive bacterium species are marked as orange circles. The genomes contains Rho or Rof are indicated as black squares.

**Fig. S8.**
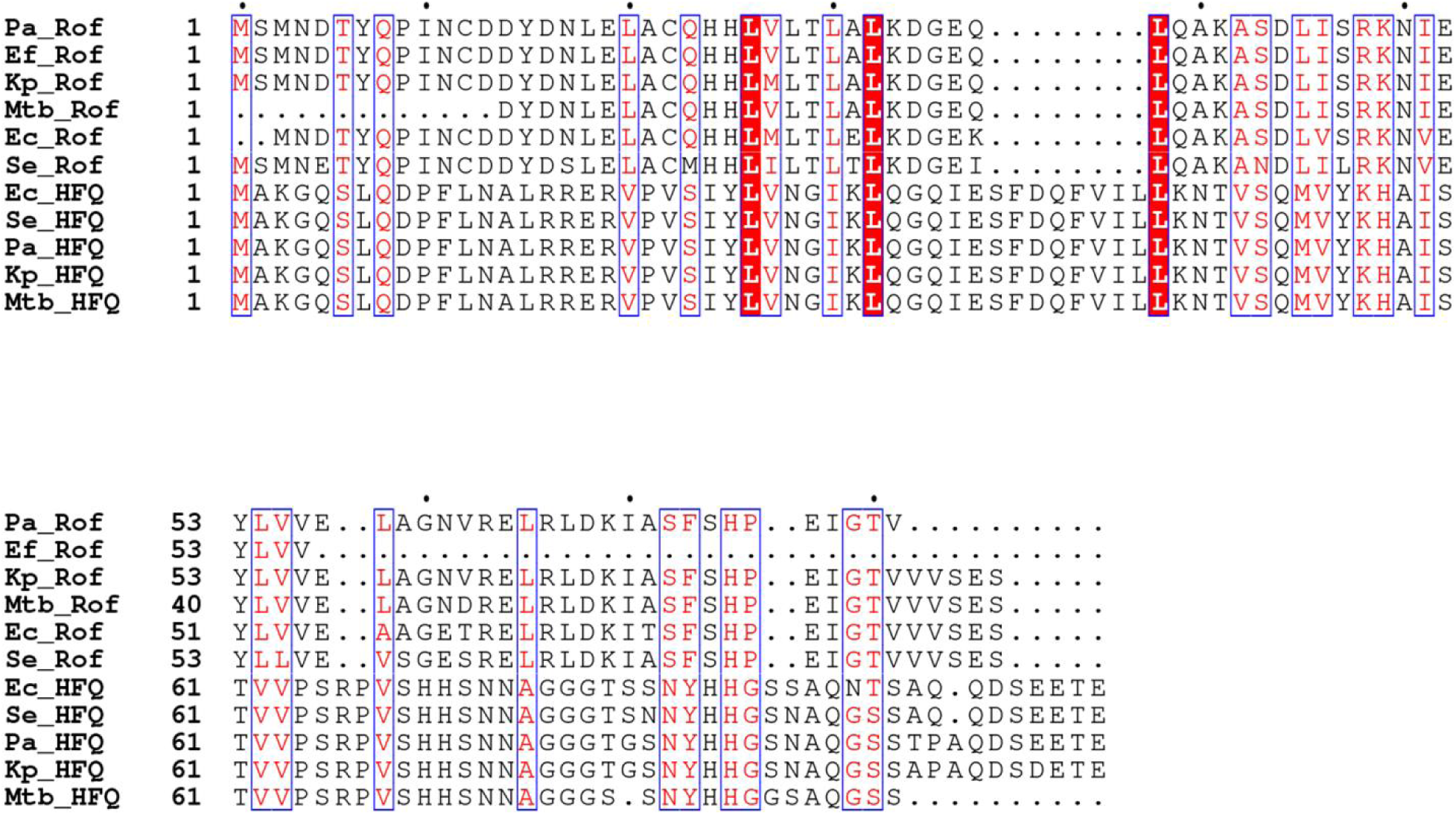
Sequence alignment of Rof and Hfq. Amino acid sequence alignment of Rof and Hfq from *Pa, Pseudomonas aeruginosa; Ef, Enterococcus faecalis; Kp, Klebsiella pneumoniae; Mtb, Mycobacterium tuberculosis; Se, Salmonella enterica; Ec, Escherichia_coli*.

**Fig. S9.**
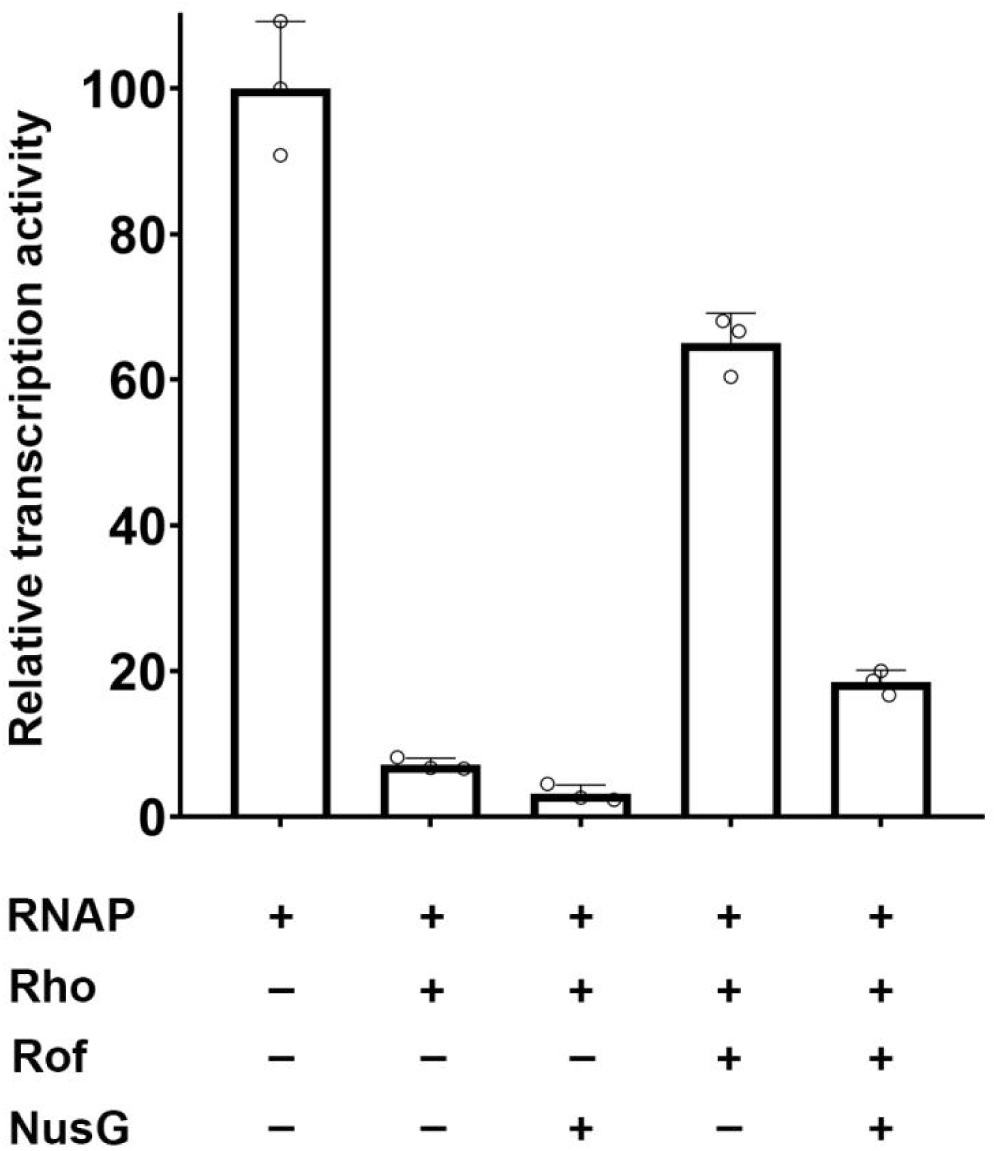
Relative *in vitro* transcription activity. Relative in *vitro* transcription activity showing NusG is functional in Rho-dependent termination. Data for *in vitro* transcription assays are means of three technical replicates. Error bars represent ± SEM of n = 3 experiments.

**Table S1:** Cryo-EM structure: E. coli Rho-Rof.

|  | **Rho-Rof**<br>**(EMDB-37342)**<br>**(PDB 8W8D)** |
| --- | --- |
| **Data collection and processing** |  |
| Magnification | 105,000 |
| Voltage (kV) | 300 |
| Electron exposure (e-/Å<sup>2</sup>) | 50 |
| Defocus range (μm) | -0.8 to -2.0 |
| Pixel size (Å) | 1.19 |
| Symmetry imposed | C1 |
| Initial particle images (no.) | 6,033 |
| Final particle images (no.) | 5,823 |
| Map resolution (Å) | 2.8 |
| FSC threshold | 0.143 |
| Map resolution range (Å) | 2.8-6.2 |
| **Refinement** |  |
| Initial model used (PDB code) | 1PV4, 1SG5 |
| Model resolution (Å) | 2.8 |

